# The Need for Sustainable Leadership in Academia – a German Case Study

**DOI:** 10.1101/2021.02.17.431650

**Authors:** Verena Haage, Linn Voss, Daniela Nguyen, Friderike Eggert

## Abstract

Academic leaders are selected based on their publication record, citation index and acquisition of third party funding. However, heading a successful research team, also requires leadership skills. Despite the clear need, leadership development has been systematically neglected in the present academic system. At the same time, growing evidence suggests that leadership styles of academic supervisors can dramatically affect the mental health of academic employees as well as drive highly skilled researchers out of academia. Here, we assessed the current state of academic leadership in the German academic system by surveying 368 participants currently employed in academia in Germany. We report that 64% of current academic leaders did not feel prepared for their current position while 86% of participants expressed their interest in leadership development programs offered by their research institutions. Our results highlight the demand for leadership development programs in German academic institutions to ensure a more efficient academic system.

## Introduction

Success in science is measured through a combination of scientific output in the form of publications in scientific journals and the acquisition of funding in order to enable further research (1, 2). As young researchers advance in their careers, they become highly trained in skills such as scientific writing so as to master publication or grant writing. The mediation of leadership skills, however, is often neglected as currently these do not contribute to the evaluation of scientific success or the appointment to faculty positions (1). Therefore, an early career researcher (ECR) may become leader of a research group based on publication record and solicitation of third party funding, but without having received sufficient training of team leadership or team development (3). A recent study focusing on leadership in academia, identified the neglect of systematic leader selection and development as one of the most pressing challenges in academic leadership, besides managing autonomy, constant change and uncertainty (4). According to the authors, academic leaders are not prepared for their demanding roles (4). Moreover, a survey including 233 professors from universities in the United Kingdom revealed that 60% indicated their research output and scholarships as the sole basis for their appointment (5).

In order to combat the so called “Peter Principle” (6) in academia, which states that *“members of an organization where promotion is based on achievement, success, and merit will eventually be promoted beyond their level of ability”*, researchers should be sufficiently trained in leadership skills for the new set of challenges and responsibilities they will face upon reaching a leading position. In fact, leadership has been considered as key to academic success (7) and combined approaches of individual as well as collective leadership have been suggested for successful research leadership (4). At the same time, growing evidence suggests that the leadership style of academic supervisors can dramatically affect mental health of academic employees, especially of PhD students (8, 9). Moreover, managing students with mental health issues can also pose enormous challenges on untrained supervisors (10), creating an unsustainable circle of insecurity and overstress due to lack of leadership skills.

Despite growing movements to advance practical and robust approaches for research assessment such as the San Francisco Declaration on Research Assessment (DORA) (11), similar movements with regard to advancing leadership skill development for academic offspring are currently rare.

Moreover, studies reflecting on the current status of research leadership are scarce. Here we surveyed 368 participants currently working in academia in Germany on their perception and experience of leadership in the German academic system, highlighting the current situation as well as the needs for change towards a more sustainable academic environment.

## Materials and Methods

The survey (Supplementary file 1) was created using the online tool SurveyPlanet and was conducted using convenience sampling with dissemination via forwarded email invitations or shared via LinkedIn, and remained open for six weeks. A pilot version of the survey was originally conducted with 4–8 doctoral researchers/PhD students of scientific research institutions in Berlin. Based on this pilot run, some questions were revised. 709 participants completed the survey. The survey was originally planned to give an international overview on the topic, since however 88.7% (629) of participants are currently working in German research institutions/academia, the subsequent analysis was focused on German academia. When asked about their highest academic degree, 7% (44) of participants stated high school diploma. According to our definition, participants should have at least a university degree to take part in a survey focused on academic leadership. For these reasons, participants who are currently fresh students but do not yet have a university degree were excluded from further analysis. Further, participants currently working in academia in Germany with at least one academic degree (585), were currently not all employed in academia, in fact 37% (217) of participants were currently working outside of academia, while 63% (368) of participants worked in academic institutions. In order to depict the current status of leadership in the German academic system, the analysis was therefore further focused on all participants currently working in the German academic system (368). All the descriptive statistics reported in this article are for these 368 respondents.

## Results

We surveyed 585 international academics currently working in Germany on their experience in leadership culture in academia, their needs for supporting leadership skill development as well as their openness towards novel leadership concepts in academia (see Methods for information on how the survey was disseminated).

Out of the surveyed German academic participants, 63% (368) are currently employed in academia, 34% (197) indicated to work outside of academia or research while 3% (20) indicated employment as scientists outside of academia. The latter two groups show experience in academia, but are currently employed in a variety of professions outside of academia; in order to reflect the current situation in academia the analysis was therefore focused on the 368 academics that are currently employed in academia. 60% (221) of participants were women, 38% (139) were men with an average age of 31 years ranging from 21 to 82 years.

The majority of participants held a PhD/MD (41%) indicating substantial experience in academic culture, followed by 38% holding a Master’s degree, while a minor part of participants held a Bachelor’s degree (21%) (Figure 1A). When asked about their current position in academia, 16% specified as Group Leaders or Professors, 19% as Postdoctoral Researchers (Post Docs), 31% as PhD Students, 14% as Research Assistants (defined as a graduate who is employed on a temporary or part-time basis to assist the university or research institution with academic research) and 20% as students (Figure 1B).

**Figure 1.**
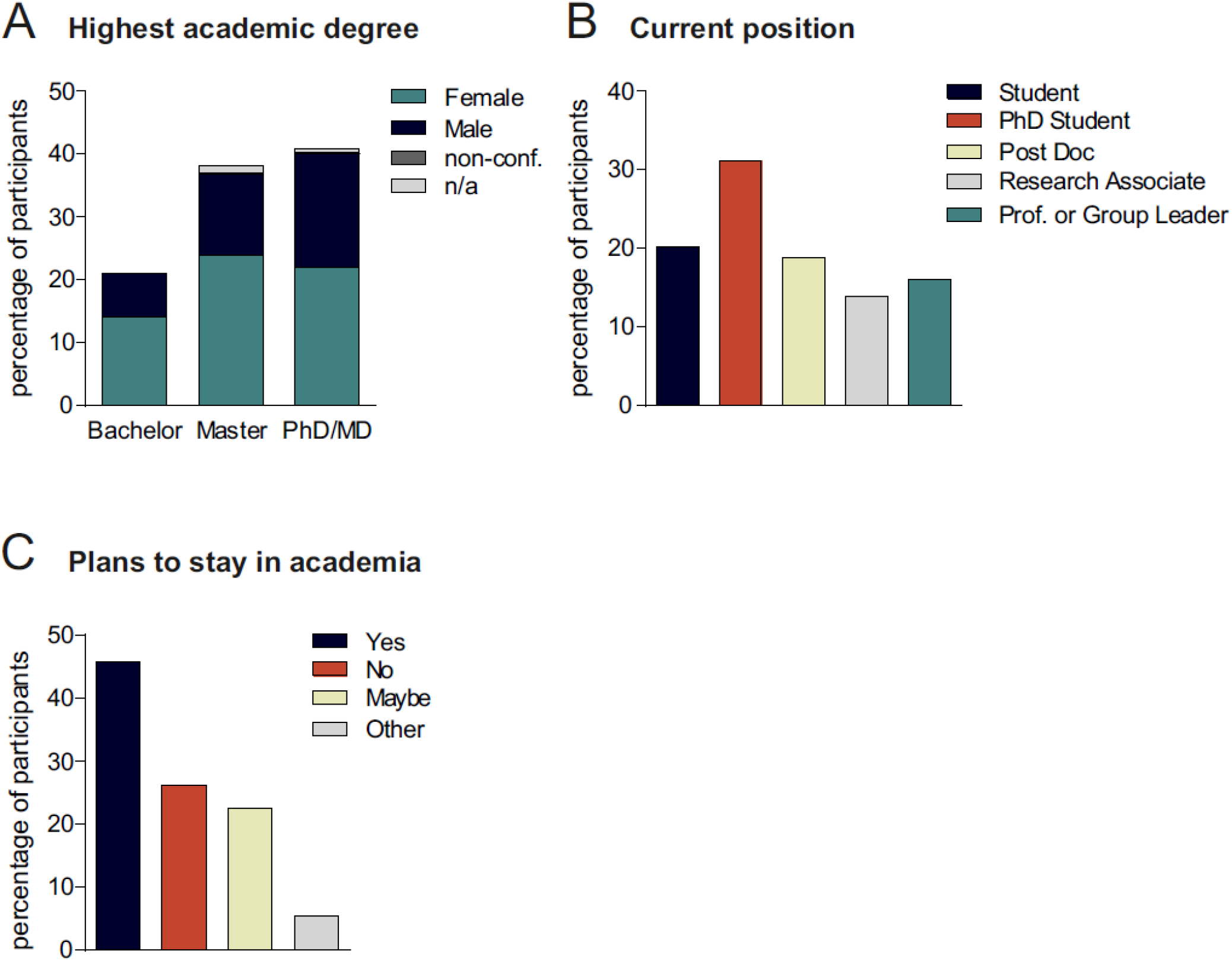
Survey demographics. **A**. Distribution of participants based on highest academic degree ranging from Bachelor, Master to PhD or MD including gender distribution. **B**. Current academic position of participants raging from Student (cyan), to PhD Student (orange), Post Doc (lime green), Research Associate (light grey) to Professor or Group Leader (turquoise). **C**. Percentage of participants planning to stay in academia (Yes; cyan), to leave academia (No; orange), is undecided (maybe; lime green), answered other (light grey). n/a: no data available.

Surveyed participants currently working in academia were further asked about their plans regarding academia. 46% indicated to plan to stay in academia (“Yes”; 55% women, 38% men), 26% (“No”; 57% women, 42% men) are planning to leave academia and 23% (“Maybe”; 72% women, 27% men) are currently undecided regarding their professional future in academia (Figure 1C). While the gender ratio was similar for staying or leaving academia (“Yes” or “No”), noticeably more women than men indicated indecisiveness (“Maybe”) regarding their future in academia. Moreover, many participants stating that they were undecided expressed their desire to stay in academia but expressed their doubts on combining a career in science with family planning, due to long working hours and short-term contracts.

When the 37% of surveyed participants that already left academia were asked about their motivation to leave, reasons were manifold; the majority, however, stated that they were concerned about poor career prospects and a lack of job security, underscoring widespread concerns of participants working in academia.

### Academics feel unprepared for leadership in academia

We further assessed whether participants working in academia feel prepared for leadership in academic environments. Out of the surveyed academic participants, 59% indicated to be currently in a leading position (53% women, 45% men) while 41% stated to be currently not in a leading position (70% women, 27% men) (Figure 2A). When asked about their plans regarding leadership, out of the 41% that are currently not in a leading position, 78% indicated to be pursuing a leading position (58% women; 38% men) while only 15% stated not to aim for a leading position (74% women; 26% men) (Figure 2B).

**Figure 2.**
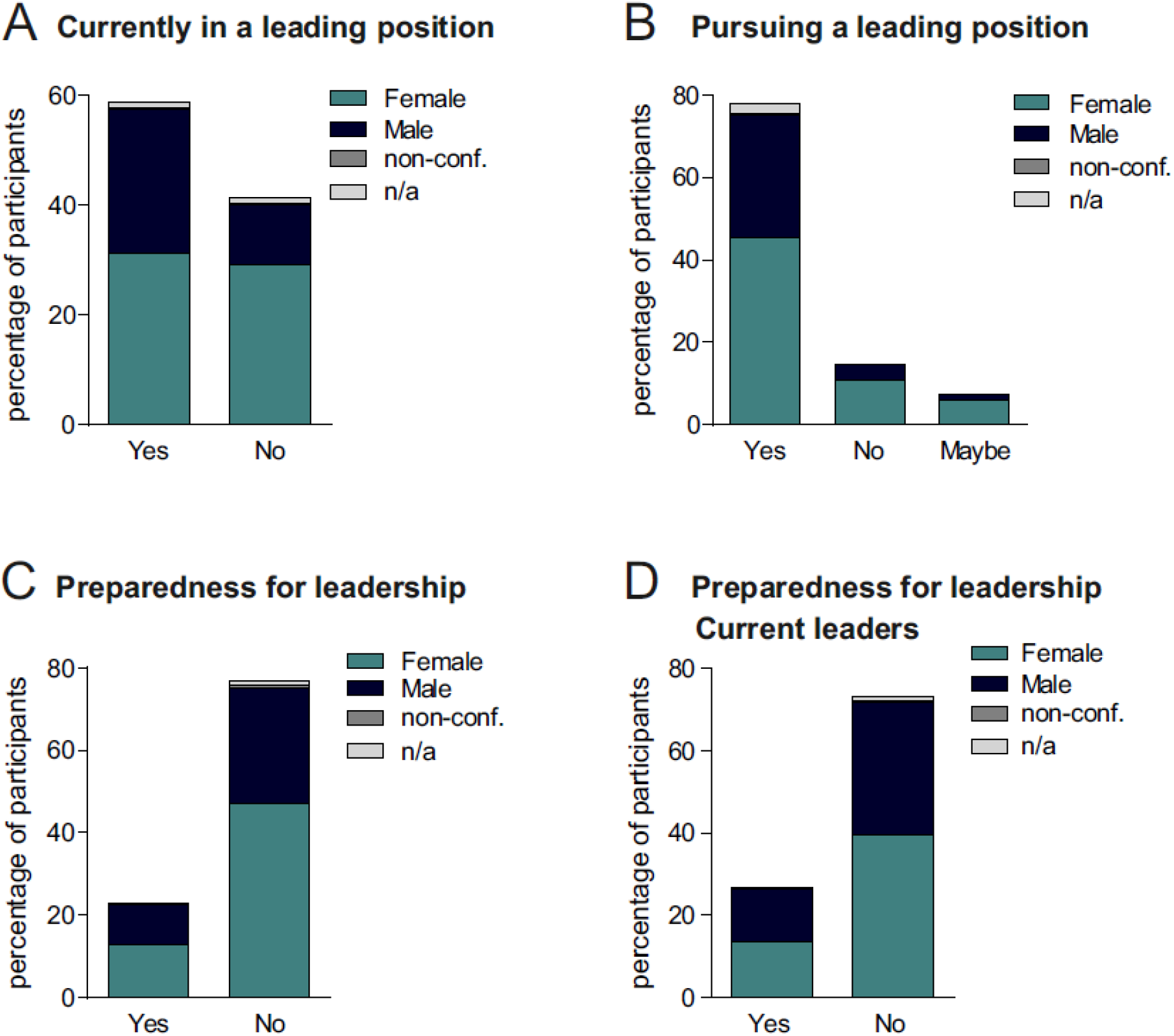
Leadership status of participants. **A**. Percentage of participants currently holding a leading position (Yes) and currently not holding a leading position (No) including gender distribution within each group. **B**. Percentage of participants pursuing (Yes), potentially pursuing (Maybe) and not pursuing (No) a leading position including gender distribution within each group. **C**. Percentage of current leaders that feel prepared (Yes) or not prepared (No) for a leading position. **D**. Percentage of current non-leaders that felt prepared (Yes) or not prepared (No) for a leading position. n/a: no data available.

Despite the majority of participants aiming for a leading position in academia, 77% of all academic participants stated that they were not well prepared for a leading position during their academic career (Figure 2C). When focusing on the current leaders in academia, 73% also stated that they did not feel well prepared for the leading position they currently hold (54% women, 44% men; Figure 2D).

When academics currently working outside of academia were asked regarding their preparedness for leadership, 51.8% of current leaders did not feel prepared for their position (Supplementary File 2).

### Academics are interested in leadership development programs and expect institutions to act

To better understand the needs of the academic community, we assessed their interest in leadership training opportunities as well as the format and conditions of such offers. 84% of participants indicated their interest in a training or coaching program supporting their leadership development (Figure 3A). When asked about the format they would prefer for leadership skill development, interests were diverse ranging from network building, to personal coaching to workshops as well as lectures or online seminars (Figure 3B).

**Figure 3.**
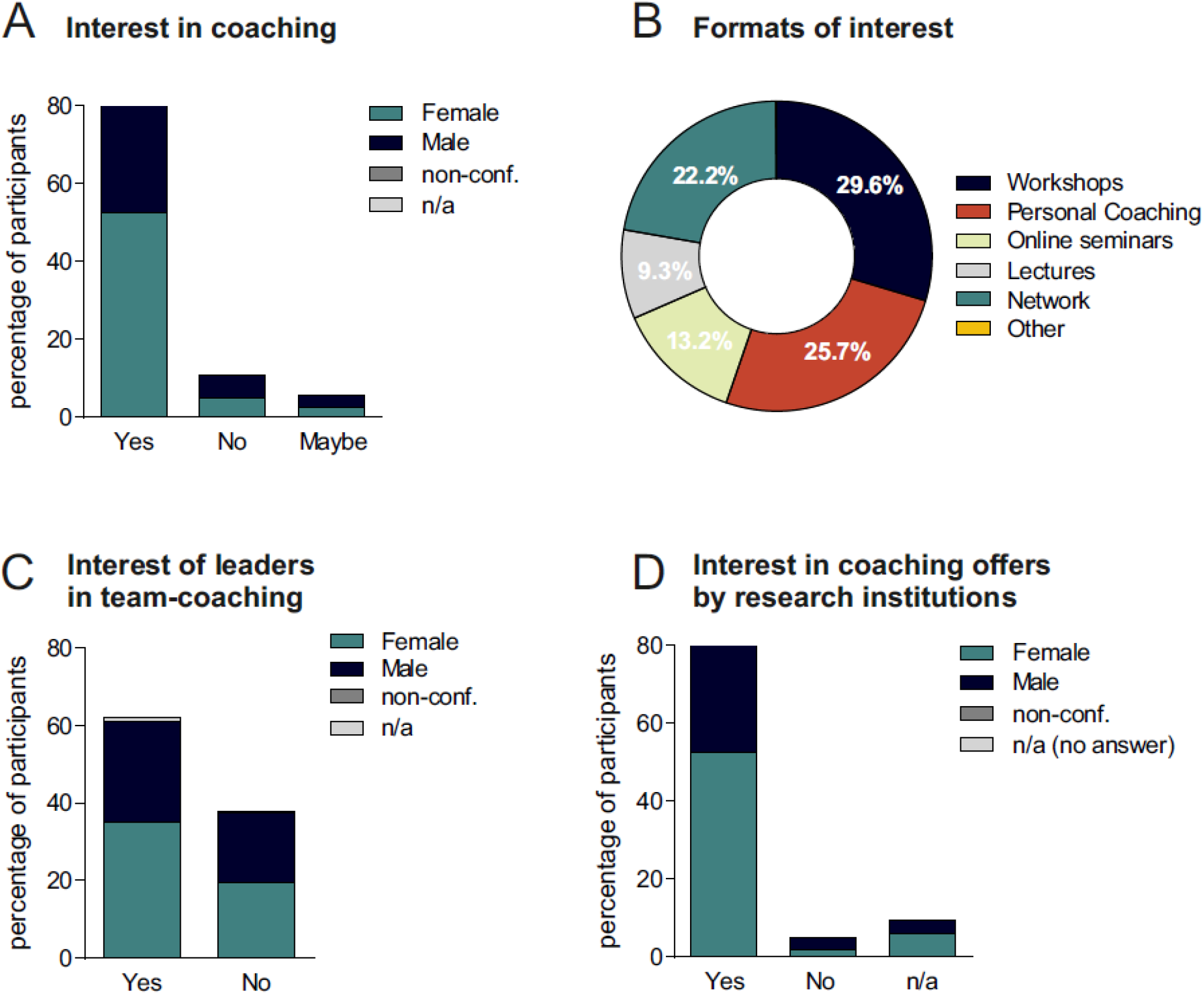
Interest in leadership training. **A**. Percentage of participants that is interested (Yes), potentially interested (Maybe), not interested (No) in coaching. **B**. Percentage of participants interested in different formats for leadership training including workshops (cyan), personal coaching (orange), online seminars (lime green), lectures (light grey), network building (turquoise), other (yellow). **C**. Percentage of current academic leaders interested (Yes) or not interested (No) in team-coaching. **D**. Percentage of participants interested (Yes) or not interested (No) in coaching offers by research institutions. n/a: no data available.

About 62% of current leaders also expressed their interest in participating in leadership development training together with their team (Figure 3C).

Having defined the needs of the academic community, we further examined the role of academic institutions in the development of leadership skills, where 86% of participants stated their interest in such offers provided by their research institutions (Figure 3D).

## Discussion

We here report a great need for leadership training programs in academia, based on data from our survey on the current state of leadership in academia in Germany. 64% of current academic leaders stated that they did not feel well prepared for the position they are currently holding while 86% of all participating academics currently employed in academia expressed their interest in leadership programs offered by their research institutions.

In current debates about academic leadership, leadership is usually defined from the perspective of a group leader or professor. From our point of view however, leadership in science starts at an earlier stage, since in order to supervise or mentor another student, a common scenario in the course of a PhD, leadership skills are already required. Here, we therefore defined leadership as an early occurring event in the course of a scientific career. Despite our definition of leadership, the majority of leaders participating in our survey were at more advanced career stages as 90% of participants held a Master&s degree, an MD or PhD degree. More advanced career stages however, indicating more time spent in academia, did not result in better preparedness for leadership, underscoring the need for leadership training at every career stage.

Due to the high number of German participants, our survey focused on German academia, reflecting the current state on leadership in academia in only one, but one of the leading countries in academic research. Research culture however might differ between countries and our data are therefore not suitable for a general statement on the current state of leadership in academia. Thus, more international studies will be required to confirm our data as well as to paint a more complete picture of the current state on academic leadership.

It is beyond debate that leadership in academia is of high complexity. Academic leaders are required to meet the interests of a spectrum of different stakeholders (12), while being held to the highest standards regarding their excellence in research and teaching (13). At the same time, academic leadership ranges across multiple levels, from an individual level, to the level of a research group to the organization (4, 14).

To date, only a few studies on the actual state of leadership in academia exist. One study that surveyed academic leaders from Chinese and European universities, reports a lack of comprehensive conceptualization of academic leadership, providing a new definition of academic leadership based from an international academic context: “an influence of one or more people with an academic profile on academic behavior, attitudes or intellectual capacity of others based on commitment and power in order to achieve managerial, structural, and institutional vision values” (15). Another study highlights the fact that many current academic leaders are actually not aware of their role in improving teaching quality at universities or learning success of their students (16). On these lines, a recent study underscores the importance of sustainable leadership practices in universities to ensure quality learning and teaching (17). According to the authors, one important component of sustainable leadership practice includes providing adequate developmental opportunities for those who are likely to become leaders of learning and teaching (17).

Similarly, our data indicate the need for leadership training for future academic leaders and at the same time their interest in such training.

Synergies from interdisciplinary collaborations, effective organization as well as diverse environments maximize the use of resources and implement a sustainable science culture in which researchers have the right framework and opportunities to focus on their projects.

One way to improve scientific leadership could be training in leadership programs that use leadership skills and working frameworks that have been applied successfully in other fields. For example, concepts such as New Work and Agility, originating in the start-up world, aim to realize an improved, innovative and creative work culture, similar to the scientific field (18, 19). These concepts are based on self-motivation and creativity, which makes them suitable for scientists, who are also strongly motivated by purpose (20, 21). Therefore, it would be plausible to incorporate them in scientific leadership training. Pioneer organizations such as the German Scholarship Organization are developing programs for scientific leaders that support the development of expertise exceeding the knowledge acquisition and scientific-work-centered education (22).

Our data indicate that the majority of current leaders in the German academic system was not prepared for their position; however, they expressed great interest in training courses that could be offered by institutions, highlighting the role of institutions in supporting the development of future scientific leaders. By investing in leadership competencies, research institutions and universities may sustainably raise the potential of academic excellence (23). Additionally, by sensitizing future academic leaders towards general obstacles facing when pursuing a scientific career such as lack of job security, power structures or imposter syndrome, reasons for many excellent researchers to leave academia, and providing support to them, institutions might contribute to sustain more researchers in academia. By promoting diversity among academic leaders, research institutions might additionally contribute to fairer and better research (24).

Some institutions have already integrated corresponding courses and are already leading by example, such as the University of Sheffield providing online resources on the development of leadership skills (25) or the Leaders Support and Development Program of the English National Institute for Health Research (NIHR) offering future-focused leadership programs for current and emerging research leaders (26). The efficacy of such programs was shown by an Australian study reporting the development of a career-development training program for early career researchers at an Australian university as well as its immediate impact on research productivity on the individual as well as organizational level (27).

## Conclusion

We found that most academics aspire to leading positions but did not feel prepared and bemoaned a lack of leadership skills in the scientific world. There might be a need to transform the science work culture from a “stick and carrot” environment where scientists work solely towards their next publication into a science enthusiasm and innovation-driven culture.

With a need for excellence in times of increasingly complex problems, leadership skills beyond mere management of teams are needed to tackle scientific questions in global collaborations. They are also needed provide role models for young researchers and provide them with future perspectives in the field of academia and a unique framework to enhance their knowledge and research skills. One answer to this question could be adopting work and leadership concepts that worked in highly innovative fields of industry such as agility to the scientific environment.

## References

1. Detsky AS. 2011. How to be a good academic leader. Journal of general internal medicine 26:88–90.

2. Fortin J-M, Currie DJ. 2013. Big Science vs. Little Science: How Scientific Impact Scales with Funding. PLOS ONE 8:e65263.

3. Lashuel HA. 2020. What about faculty? Elife 9.

4. Braun SaP, C. and Frey, D. and Knipfer, K.. 2016. Leadership in academia: individual and collective approaches to the quest for creativity and innovation. Leadership lessons from compelling contexts Bingley: Emerald: pp. 349–365.

5. Macfarlane B. 2011. Professors as intellectual leaders: formation, identity and role. Studies in Higher Education 36:57 –73.

6. Peter LJaH, Raymond.. 1969. The Peter Principle.

7. Bryman A. 2007. Effective leadership in higher education: a literature review. Studies in Higher Education 32:693–710.

8. Levecque K, Anseel F, De Beuckelaer A, Van der Heyden J, Gisle L. 2017. Work organization and mental health problems in PhD students. Research Policy 46:868–879.

9. Christian K, Johnstone C, Larkins JA, Wright W, Doran MR. 2021. A survey of early-career researchers in Australia. Elife 10.

10. Loissel E. 2019. A question of support. Elife 8.

11. https://sfdora.org/. Accessed 17.01.2021.

12. Milliken J. 1998. The Cult of Academic Leadership.. Higher Education in Europe 23(4):505–515.

13. Corlett JA. 2005. The good professor. Journal of Academic Ethics 3(1):27–54.

14. Bolden R, Petrov, G., & Gosling, J. 2009. Distributed leadership in higher education: What does it accomplish? Leadership 5(3):299–310.

15. Dinh NBK, Caliskan A, Zhu C. 2020. Academic leadership: Perceptions of academic leaders and staff in diverse contexts. Educational Management Administration & Leadership doi:10.1177/1741143220921192.

16. Alenoush Saroyan DG, Engida Gebre. 2011. Understanding academic leadership, abstr Annual meeting of the American Educational Research Association,

17. Bosanquet A, Cameron A, Marshall S, Orrell J. 2021. Ensuring Sustainable Leadership for Quality Learning and Teaching.

18. Nafei W. 2016. Organizational Agility: The Key to Organizational Success. International Journal of Business and Management 11:296.

19. Sherehiy B, Karwowski W. 2014. The relationship between work organization and workforce agility in small manufacturing enterprises. International Journal of Industrial Ergonomics 44:466–473.

20. McGee Ebony O, White Devin T, Jenkins Akailah T, Houston S, Bentley Lydia C, Smith William J, Robinson William H. 2016. Black engineering students’ motivation for PhD attainment: passion plus purpose. Journal for Multicultural Education 10:167–193.

21. Johnson BB, Dieckmann NF. 2020. Americans’ views of scientists’ motivations for scientific work. Public Underst Sci 29:2–20.

22. https://gsonet.org. Accessed

23. Riccio S. 2010. Talent Management in Higher Education: Developing Emerging Leaders Within the Administration at Private Colleges and Universities.

24. Aguirre A. 2017. Diversity and Leadership in Higher Education, p 1–8. In Teixeira P, Shin JC (ed), Encyclopedia of International Higher Education Systems and Institutions doi:10.1007/978-94-017-9553-1_533-1. Springer Netherlands, Dordrecht.

25. https://www.sheffield.ac.uk/rs/ecr/training/psrl. Accessed

26. https://www.nihr.ac.uk/explore-nihr/academy-programmes/nihr-leaders-support-and-development-programme.

27. Browning L, Thompson K, Dawson D. 2014. Developing future research leaders: Designing early career researcher programs to enhance track record. International Journal for Researcher Development 5:123–134.

