## Supplementary Files for "The Need for Sustainable Leadership in Academia – a German Case Study"

**Supplementary File 1: Survey Questions**

1. Which gender identity do you identify most with? (multiple choice)
    - Female
    - Male
    - Gender-variant
    - non-conforming
    - not listed
    - prefer not to answer

2. What is your age? (Essay)

3. What is your highest degree? (multiple choice)
    - High school diploma
    - Bachelor´s degree
    - Master´s degree
    - PhD
    - MD

4. What is your current position? (Essay)

5. A. If you are working in academia: Are you planning to stay in academia after completing your current position? (Essay)

5. B. If you left academia: What was your motivation to leave? what have you learnt from it? (Essay)

6. Are you currently in a leading position? (multiple choice)
*(from supervision of students to running a small team, also outside of academia)*
    - Yes
    - No
    - Comment (Essay)

7. If currently you are not in a leading position, are you pursuing a leading position? (multiple choice)

- Yes
- No
- Comment

8. Do you feel that you were prepared/trained/briefed in the course of your career or by your institution for being in a leading position? (multiple choice)

- Yes
- No

9. Would you be interested in a training/coaching service/program supporting you in building your personal leadership and team structures (with the goal to provide you with more time for effective research/work)? (multiple choice)

- Yes
- No

10. What would you expect from such a service/program? (multiple choice)

- Workshops
- In-person coaching
- Webinars
- Lectures
- Building a network
- Other

11. In case you are in a leading position already, would you be interested in using a leadership/team organizational workshop with your team/research group? (multiple choice)

- Yes
- No
- Comment (Essay)

12. Would you like your institution to offer such a service or at least provide you with the information and offer of such? (multiple choice)

- Yes
- No
- Comment (Essay)

**Supplementary File 2**

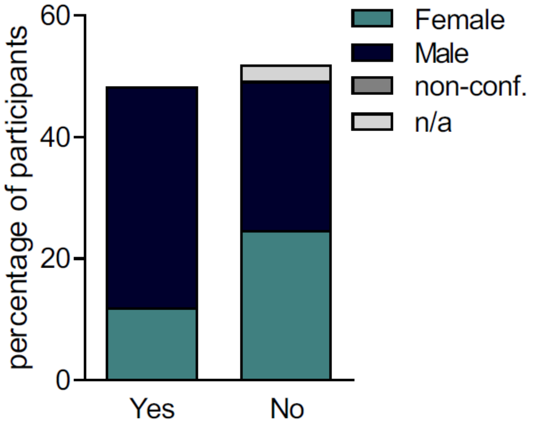


**Supplementary Figure 1. Preparedness of leaders working outside of academia.** Percentage of current leaders outside of academia that feel prepared (Yes) or not prepared (No) for a leading position.
